# Maternal antibodies provide partial protection from postnatal Zika viremia in nonhuman primates

**DOI:** 10.1101/612754

**Authors:** Nicholas J Maness, Blake Schouest, Anil Singapuri, Maria Dennis, Margaret H. Gilbert, Rudolf P. Bohm, Faith Schiro, Pyone P. Aye, Kate Baker, Koen K. A. Van Rompay, Andrew A. Lackner, Myrna C. Bonaldo, Robert V. Blair, Sallie R. Permar, Lark L. Coffey, Antonito T. Panganiban, Diogo Magnani

## Abstract

Zika virus (ZIKV) will remain a public health threat until effective vaccines and therapeutics are made available in the hardest hit areas of the world. Recent data in a nonhuman primate model showed that infants postnatally infected with ZIKV were acutely susceptible to high viremia and neurological damage, suggesting the window of vulnerability extends beyond gestation. We addressed the susceptibility of two infant rhesus macaques born healthy to dams infected with Zika virus during pregnancy. Passively acquired neutralizing antibody titers dropped below detection limits between 2 and 3 months of age, while binding, possibly non-neutralizing antibodies remained detectable until viral infection at 5 months of age. Post-infection acute serum viremia was substantially reduced relative to adults infected with the same dose of the same stock of a Brazilian isolate of ZIKV (n=11 pregnant females) and another stock of the same isolate (n=4 males and 4 non-pregnant females). Virus was never detected in cerebrospinal fluid nor in neural tissues at necropsy two weeks after infection, suggesting reduced viral burden relative to adults and published data from infants. However, viral RNA was detected in lymph nodes, confirming some tissue dissemination. Though protection was not absolute, our data suggest infants born healthy to infected mothers may harbor a modest but important level of protection from postnatally acquired ZIKV for several months after birth, an encouraging result given the potentially severe infection outcomes of this population.

## Introduction

Zika virus (ZIKV) emerged in Brazil in 2015 and maternal infection during pregnancy was astutely correlated with an increase in newborns with microcephaly [1, 2], a profound developmental defect that results in infants with reduced brain size and cognitive capacity. ZIKV was initially discovered in 1947 in the Zika forest of Uganda during surveillance for yellow fever virus [3]. Soon thereafter, it became clear that human infections with ZIKV in that region were not uncommon [4, 5] but disease associated with infection appeared to be minor and ZIKV became something of an afterthought. The emergence in Brazil and its association with both major and, more recently, less severe neurological consequences in congenitally-infected newborns [6], collectively called “congenital Zika syndrome”, rapidly changed that perception. A profound research effort was subsequently launched to understand mechanisms of pathogenesis [7, 8], identify cellular receptors [9–12] and targets [9, 12–16], to develop animal models [17–23], and to develop and test vaccines [24–29] and therapeutics [23, 30].

Human brain development continues well after birth [31] so it stands to reason that the risk of ZIKV associated neurological disease may extend for an unknown period of time after birth. Indeed, a recent study in nonhuman primate infants showed high peak viral loads and dissemination into multiple brain regions at two weeks post-infection and quantifiable neurological defects and cognitive impairment in infants infected in the first few months of life [32].

Given the high incidence of ZIKV infection in several South and Central American countries during the height of the ZIKV epidemic, it is likely that a large number of babies without congenital infection or disease sequelae were born to infected mothers. The vulnerability of these infants to newly acquired infection after birth has not been addressed. A recent macaque study showed that fetal infection after subcutaneous inoculation of dams with ZIKV was efficient, with four of four fetuses showing evidence of infection [33]. Since not all infants exhibit detectable ZIKV disease when born to infected mothers, it remains unclear whether these infants remain uninfected and/or unaffected due to pre-existing passively acquired maternal antibodies, if they mount their own de novo anti-ZIKV immune responses in utero or soon after birth, or if infection can be limited and apathogenic for an unknown reason. Addressing these issues will be key to addressing the susceptibility of newborns to postnatal ZIKV infection in areas with endemic for ZIKV transmission. During this study, we monitored antiviral antibody responses after birth in two infant macaques born to ZIKV infected dams and assessed the level of protection these responses might provide against postnatal infection. Upon infection at five months of age, both infants showed only modest levels of peripheral viremia and no virus detected in neurological tissues. These data suggest that being born to a ZIKV infected mother may confer a small but important level of immunity to postnatal infection.

## Materials and Methods

### Cohort

All macaques used in this study were housed at the Tulane National Primate Research Center (TNPRC), which is fully accredited by AAALAC (Association for the Assessment and Accreditation of Laboratory Animal Care) International, Animal Welfare Assurance No. A3180-01. Animals were cared for in accordance with the NRC Guide for the Care and Use of Laboratory Animals and the Animal Welfare Act. Animal experiments were approved by the Institutional Animal Care and Use Committee (IACUC) of Tulane University (protocol P0336). Two adult female, purpose bred Indian rhesus macaques, were identified as pregnant and subsequently assigned to the study. These macaques were infected with ZIKV during early third trimester. Infants were delivered via caesarian section at approximately gestational day 155 (full term) and housed in a primate nursery until 5 months of age and then infected with ZIKV subcutaneously with 10^4 PFU of a Brazilian isolate (Rio-U1/2016 GenBank KU926309), which was passage twice in Vero cells post-virus isolation. The animals were euthanized fourteen days later.

### Viral load measurements

Viral RNA was amplified and quantified as described previously [23]. Briefly, RNA was manually extracted from fluid samples (CSF or blood serum) using the High Pure Viral RNA Kit (Roche). RNA was then subjected to reverse transcription and quantitative PCR using primers and a fluorescently conjugated probe on an Applied Biosystems 7900 instrument.

### Plaque Reduction Neutralization Test (PRNT) 80 measurements

Neutralizing antibody quantification by plaque reduction neutralization test (PRNT) endpoint 80% PRNT titers were determined in infant macaque plasma, where each sample was tested in duplicate. Plasma samples were heated to 56°C for 30 minutes to inactivate complement, serially 2-fold diluted starting at 1:10 (1:20 final virus:plasma dilution) in 150 µl Dulbecco’s Modified Eagle Medium (DMEM) with 2% fetal bovine serum, and then incubated for 1 hour at 37°C with approximately 100 plaque forming units of a 2015 Brazilian ZIKV strain (SPH2015, GenBank accession number: KU321639.1) from a third Vero cell passage. After 1 hour, virus-antibody or virus-only mixtures were overlaid on confluent African Green Monkey Kidney (Vero) cell monolayers and incubated for 1 hour with rocking every 15 minutes. The plaques developed under 0.5% agar overlays in DMEM were counted after 7 days under crystal violet staining. Dilutions of plasma that caused a >80% reduction in the number of plaques, as compared with negative controls (DMEM only), were considered positive. The reciprocal of the highest dilution of plasma (represented as the mean final virus-serum dilution from both replicates) that inhibited at least 80% of plaques is reported as the antibody titer.

### Detection of ZIKV-specific IgG in rhesus plasma

High-binding 96-well ELISA plates (Greiner; Monroe, NC) were coated with 40 ng/well of 4G2 monoclonal antibody, produced in a mouse hybridoma cell line (D1-4G2-4-15,ATCC; Manassas, VA), diluted to 0.8 ng/uL in 0.1M carbonate buffer (pH 9.6) and incubated overnight at 4°C. Plates were blocked in 1X Tris-buffered saline containing 0.05% Tween-20 and 5% normal goat serum for 1 hour at 37°C, followed by an incubation with diluted ZIKV (strain PRVABC59, BEI; Manassas, VA) for 1 hour at 37°C. Optimal virus dilution was determined by whole virion ELISA (WVE) and a 1:5 dilution was used in these assays. Plasma samples were tested at a dilution of 1:12.5-204,800 in serial 4-fold dilutions and incubated for 1 hour at 37°C, along with a ZIKV-specific monoclonal antibody, H24 (10 ug/mL), isolated from a ZIKV-infected rhesus macaque. Horseradish peroxidase (HRP)-conjugated mouse anti-monkey IgG secondary antibody (Southern BioTech; Birmingham, AL) was used at a 1:4,000 dilution and incubated at 37°C for 1 hour, followed by the addition of SureBlue Reserve TMB Substrate (KPL; Gaithersburg, MD). Reactions were stopped by Stop Solution (KPL; Gaithersburg, MD) after a 7-minute incubation per plate in the dark. Optical density (OD) was detected at 450 nm on a Victor X Multilabel plate reader (PerkinElmer; Waltham, MA). Binding was considered detectable if the sample OD value at the lowest dilution was greater than that of the Background OD, defined as the OD value of the negative control at the lowest dilution plus 2 x standard deviations (SD). For samples considered positive, their OD values for the serial dilution were entered into Prism v8 (GraphPad Software; San Diego, CA) to determine the 50% effective dilution (ED_50_). The ED_50_ was calculated by first transforming the x-axis values, the dilution series 12.5-204,800 4F, into Log_10_. The transformed data was then analyzed using a sigmoidal dose-response nonlinear regression model. Any sample considered negative was assigned an ED_50_ of 12.5, the lowest dilution tested, because ED_50_ cannot be accurately calculated below the lowest dilution tested. Zika-specific IgG binding was reported in Log_10_ ED_50_.

### Behavioral observations

We employed a battery of age-appropriate behavioral tests that are designed for use in infant nonhuman primates. These tests were performed to identify any effects prenatal exposure to ZIKV. Both infants received neurobehavioral tests modelled upon testing tools used for human infants [34, 35] and adapted for use in nonhuman primates [36]. Tests were administered every two weeks, from 14 days of age until euthanasia at 20 (F10) or 21 weeks (F09). Each infant’s scores were compared descriptively against the mean and standard deviation across seven control animals reared in the same fashion and tested by the same behavioral technician. Data for control animals were available at three time points.

During the first month of life, a Neonatal Behavioral Assessment (NBA) tool was employed. Scores derived from 47 testing elements grouped for analysis into four categories, clustered by previous factor analysis [36]: orientation, state control, motor maturity, and activity. After infants reached 30 days of age, Bayley tests were administered. Scores from 48 testing elements were grouped for analysis into three categories, cognition, motor abilities, and temperament state.

## Results

### Infants born healthy with no evidence of viral infection

Both infants enrolled in this study were born via caesarean section at full term to dams infected in the third trimester as part of a previous study [23]. At the time of caesarean section, both dams had cleared serum virus but one dam exhibited a spike of amniotic fluid virus that remained detectable at the time of caesarean section (Figure 1A). However, at birth, neither infant showed evidence of infection as measured in blood or cerebrospinal fluid (CSF) (Figure 1B).

**Figure 1.**
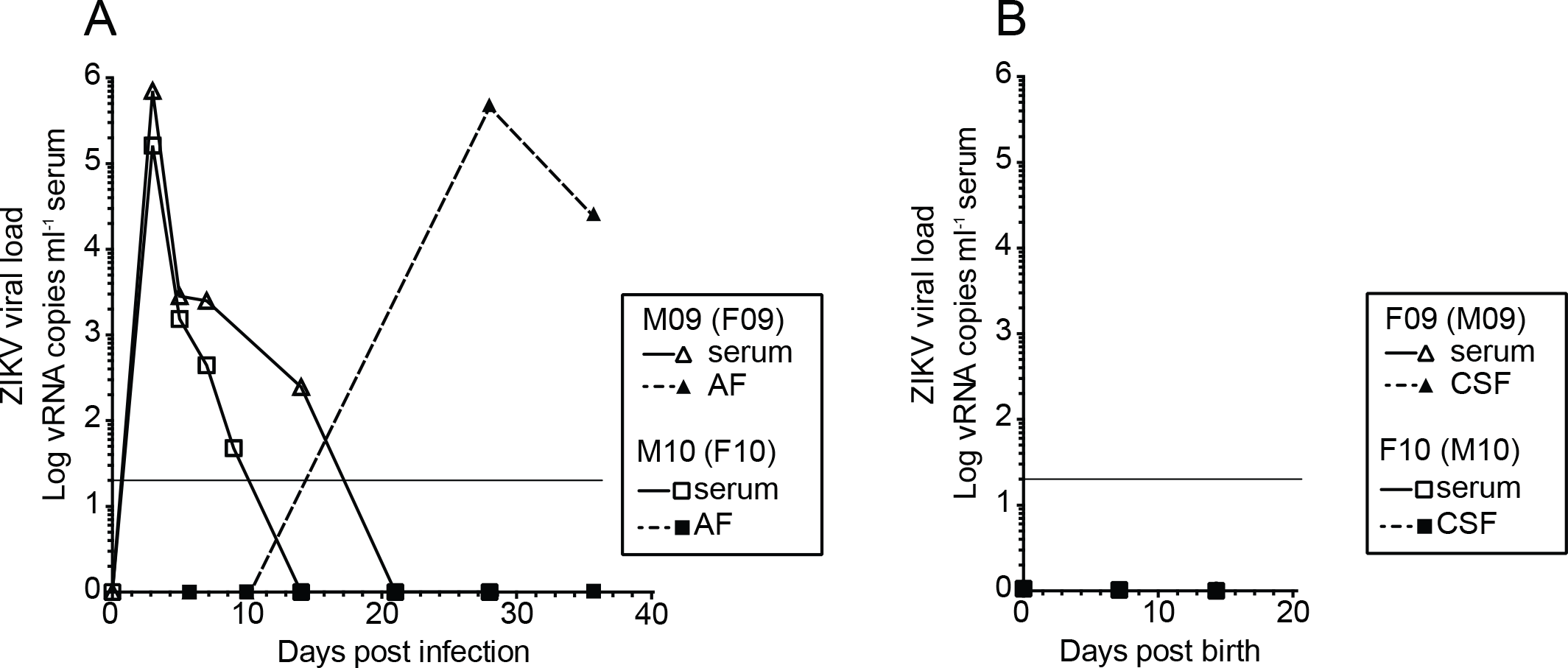
Viral dynamics in blood and amniotic fluid in two female macaques (dams of the infants in this study) (A). Each animal was inoculated with ZIKV during early third trimester and monitored for infection until giving birth via cesarean section at full term (approximately gestational day 155). Neither infant had detectable virus in serum or CSF during the first two weeks of life (B). The limit of detection of approximately 15 copies per milliliter of plasma is shown as a horizontal line in both panels.

Despite no direct evidence of infection in the infants, we next assessed the possibility that infection had occurred *in utero* and induced neurological deficits. To do this, we used a battery of defined testing parameters to compare the infants with seven control infants raised in the same manner and tested by the same technician. At 15 days of age, both infants showed slightly elevated levels of state control, motor maturity and activity while one infant did not attend to the orientation test (Figure 2A). At 16 and 20 weeks of age, both infants showed levels of cognitive abilities that were somewhat elevated while motor development was normal. The temperament of both infants was markedly calmer than controls (Figure 2B). The small sample size negated any meaningful statistical analysis. These behavioral observations were exploratory and were designed to identify any overt abnormalities that might be explored in future studies and none were noted.

**Figure 2.**
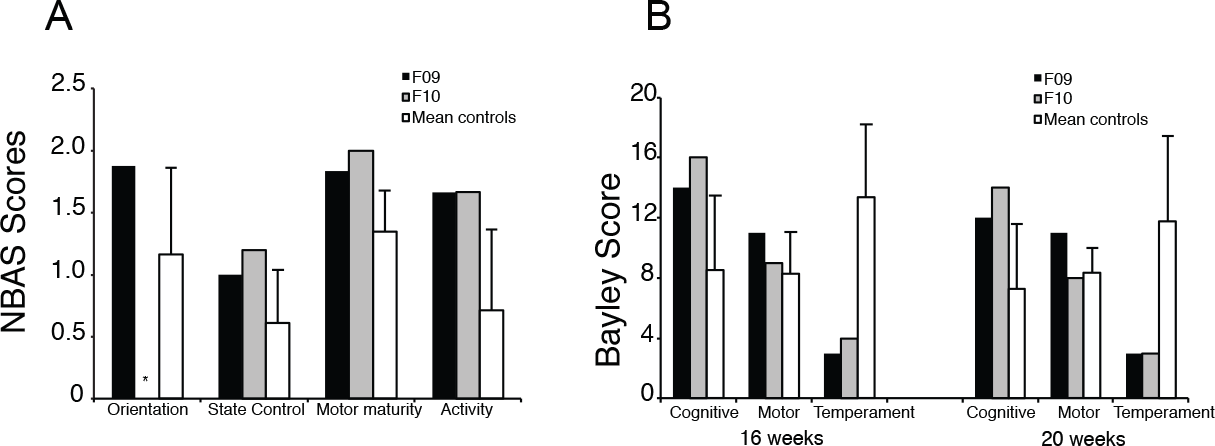
Neurobehavioral scores for F09 and F10 and 7 controls (mean+SD). Both infants were assessed for behavioral abnormalities and test scores were measured at 15 days of age (A), as described in the text. F10 was not attentive during the test of orientation so was not assessed (indicated by an asterisk *). Each infant was likewise assessed for cognitive, motor, and temperament at sixteen and twenty weeks of age (B). The only variable that showed significant differences from control animals was temperament, at both time points.

### Viral dynamics in the infants

At approximately five months of age (148 days for F10, 155 days for F09), we inoculated both infants with the same dose (10^4 PFU), via the same subcutaneous route, of the same Brazilian isolate of ZIKV that their dams had been infected with. Peak viral load in infant F09 was approximately 20,000 viral RNA copies per milliliter of plasma, which was rapidly and completely cleared by day 5 post infection. F09 was the only animal in our studies to clear blood viral RNA prior to day 5. In infant F10, the viral load remained below 1,000 copies per milliliter but remained detectable until day 7 (Figure 3A). These acute viral loads contrast with those of 11 pregnant females infected with the same stock of the same strain of the virus at the same dose and route (Figure 3B) as well as four non-pregnant females (Figure 3C) and four adult males (Figure 3D) infected with a separate stock of the same dose and strain of the virus. Area under the curve (AUC) analyses showed that F09 had a total viremia lower than all other animals in our previous studies with the exception of a single pregnant female that had a slightly lower peak viremia and cleared virus from blood far earlier than was typical for our pregnant animals. F10 had total viremia far lower than any animal in any cohort tested at our facilities (Figure 3E). At necropsy, we performed RT PCR for ZIKV RNA on serum, CSF, multiple brain regions (frontal cortex, parietal lobe, occipital lobe, temporal lobe, brain stem, optic nerve, cerebellum, choroid plexus, and subcortical white matter), and axillary lymph nodes and virus was detected only in the axillary lymph in both animals (Figure 3F). These data contrast sharply from a recent study that found infants born to healthy dams and infected postnatally showed viral loads that peaked between 10^6 and 10^7 viral copies per milliliter, which is approximately one log higher than that demonstrated by the adults and where viral RNA was detected in several neurological sites two weeks post infection [32]. Neither infant in this study showed signs of potential virus-induced pathology at necropsy. F10 harbored a choroid plexus cyst that resulted in unilateral hydrocephalus in the brain, but such cysts are common, are generally considered of little consequence and are not likely viral in origin.

**Figure 3.**
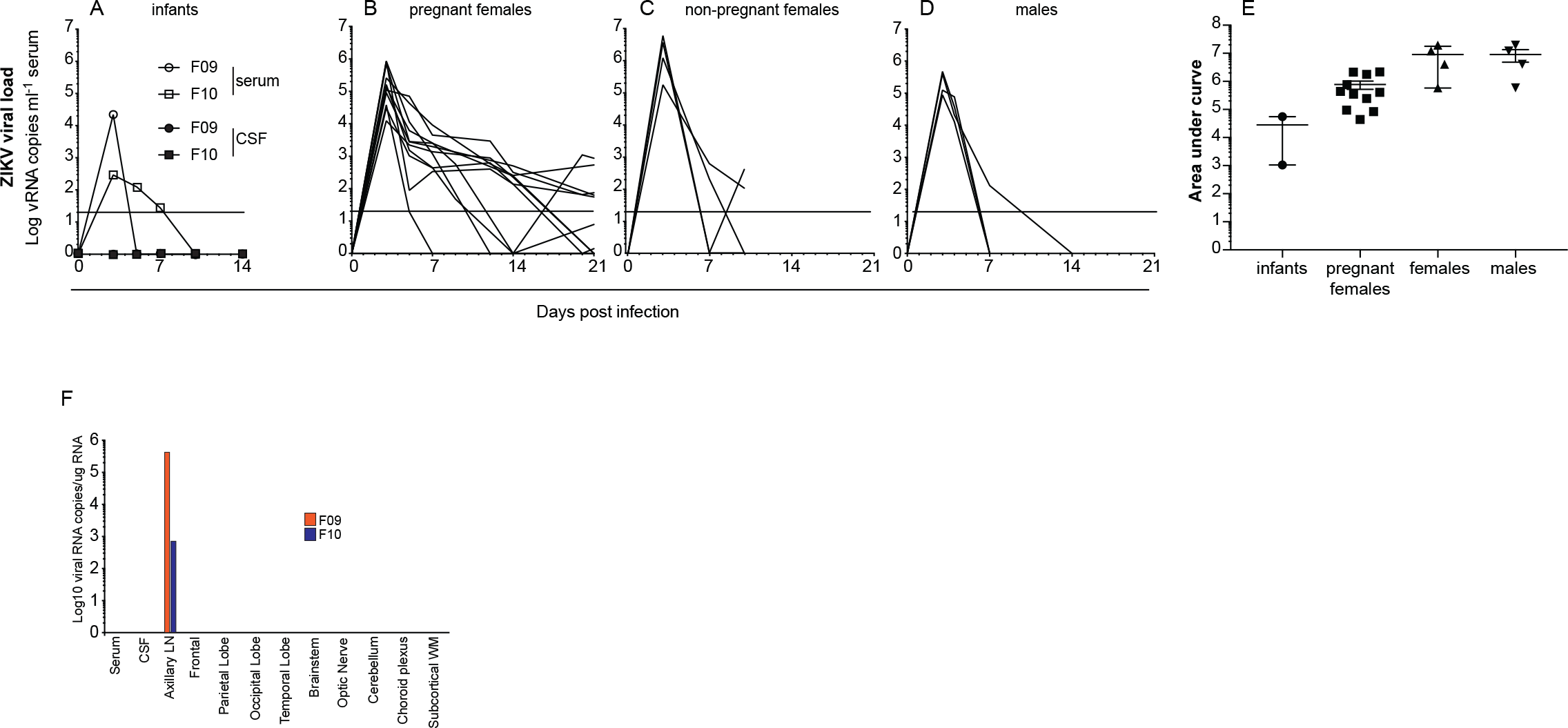
Viral dynamics in the infants after infection. Viral loads (serum and CSF) were assessed at days 0, 3, 5, 7, 10, and at necropsy on day 14 in both infants (A). For comparison, viral loads are shown for eleven pregnant females (B) infected with the same dose and route of the same stock of virus, as well as four adult non-pregnant females (C), and four adult males (D) infected with the same dose and route of a separate stock of the same isolate of ZIKV, which was passaged an additional time in Vero cells. The limit of detection of approximately 15 copies per milliliter of plasma is shown as a horizontal line. Viremia remained detectable beyond 21 days in several pregnant females but these values are cut from panel (B) for clarity. Area under the curve analysis (E) showed lower total viremia in our infants relative to nearly all other animals in our studies. Data from the pregnant females includes all time points with viremia, including beyond day 21. At necropsy, the presence of ZIKV viral RNA was assessed by qRT PCR from blood, CSF, and several brain sections as well as a lymph node from each animal (F).

### Antibody responses

We next examined humoral responses in the infants to see if they might explain the strikingly low viral loads. We used a plaque reduction neutralization test (PRNT) to assess neutralizing antibodies in serum after birth and after infection in both infants. Both showed detectable levels of neutralization at birth, which quickly waned below the limit of detection by 2 to 3 months. Neutralizing antibodies reemerged after infection and continued to rise until euthanasia at 2 weeks post infection (Figure 4A). To measure binding antibodies, we employed a whole virion ELISA assay using plasma samples collected throughout the infants’ lives both before and after infection. Binding IgG titers decreased between birth and 3-4 months of age, consistent with the expected kinetics of passively-transferred maternal IgG, but remained detectable until viral inoculation at five months, and then rose after infection, similar to the neutralizing antibody titers (Figure 4B).

**Figure 4.**
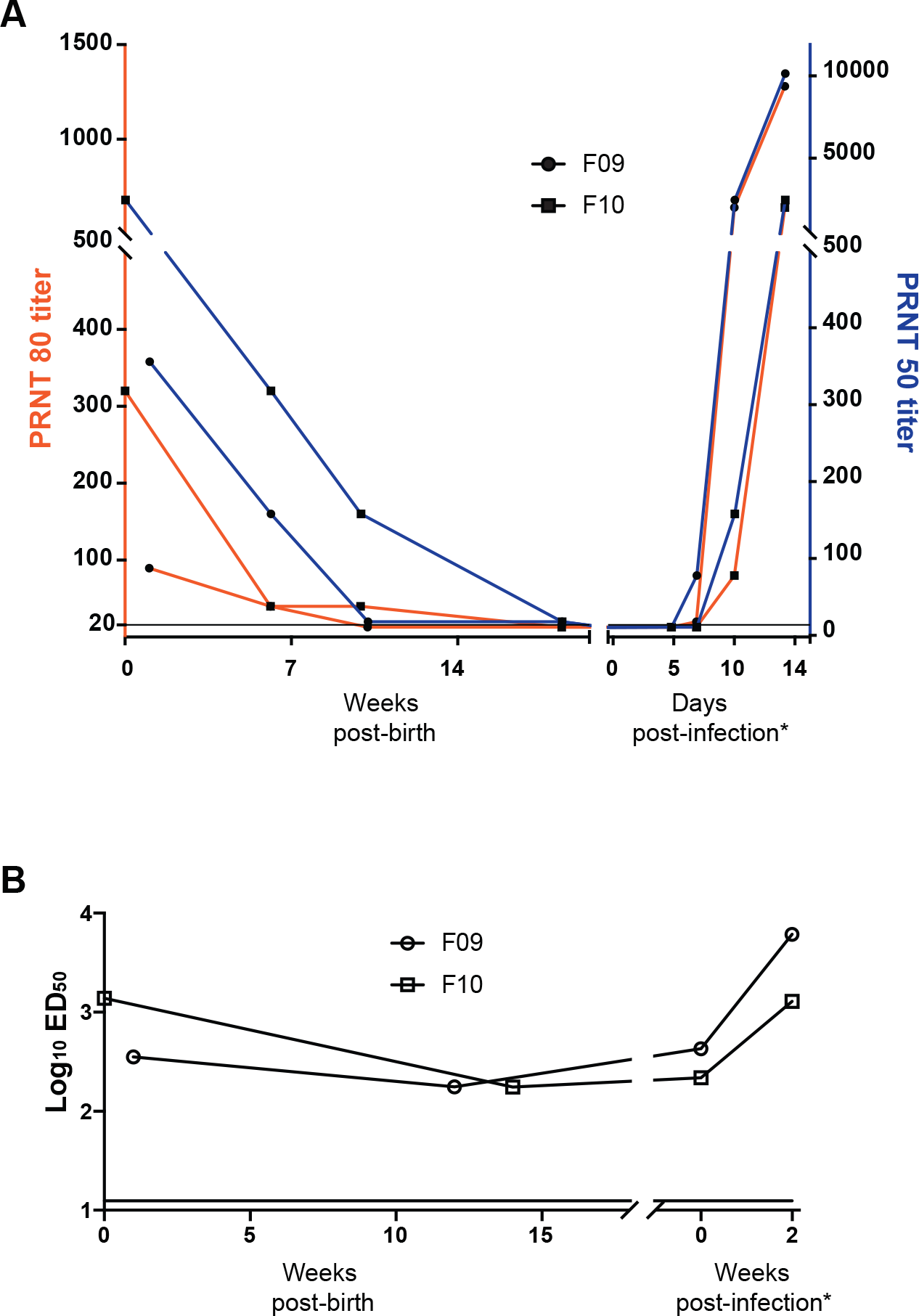
ZIKV-specific antibody responses. Neutralizing antibodies were tested using a Plaque reduction neutralization test (PRNT) (A). PRNT 80% values of plasma from both infants born to ZIKV-infected mothers over the first five months of life declined to undetectable levels by 4 months, prior to infection with ZIKV. Plasma samples with PRNT 80% titers of <20 are reported at 20. Each sample was tested in duplicate; the average titer is shown. Anti-ZIKV binding antibodies were assessed using a whole virion ELISA (WVE) test (B). Binding antibodies remained detectable until infection and expanded after infection. Each sample was tested in duplicate and the average titer is shown. The lower limit of detection in this assay was determined to be 12.5 (horizontal line).

## Discussion

ZIKV reemerged in 2015 in South America and was quickly correlated with an increase in infants born with profound neurological defects [1, 2]. It’s not entirely clear what fraction of pregnant women that become infected with ZIKV during pregnancy transmit the virus to their developing fetus but consequences to the fetus can range from mild to severe. Additionally, nonhuman primate studies suggest that infants may remain susceptible to ZIKV induced disease even if infected after birth [32], but the frequency and disease severity postnatal ZIKV infections in human infants remains to be fully examined [37, 38].

Given the high incidence of ZIKV in several countries during the height of the outbreak [39], the relative abundance of the most competent vector for ZIKV transmission, the mosquito *Aedes aegypti*, particularly in urban environments (Centers for Disease Control [CDC], 2017), it’s likely that many infants born healthy to infected mothers are themselves exposed to the virus via mosquito bite after birth. It is not clear if passively acquired maternal antibodies against ZIKV can offer some level of protection and for what period of time after birth. It is also not clear if infants exposed to the virus in utero, but born healthy and seemingly uninfected, may themselves have mounted de novo immune responses against the virus in utero. Few examples of adaptive immune responses induced in utero are described. Functional, malaria-specific T cell responses have been detected in fetuses [40], and infants are routinely vaccinated against hepatitis B virus within twenty four hours of birth, which has dramatically reduced the frequency of infant infection [41], suggesting a high level of immune competence very early in life. In the context of ZIKV, macaque data suggest that vertical transmission is quite common [33] but, to date, there is no data suggesting these infants mount antiviral adaptive immune responses. In contrast, passively acquired maternal antibodies are fairly well described. Their magnitude, transmission efficiency in utero, and decay kinetics after birth have been described in the context of infection with and vaccination against several pathogens [42, 43]. To date, no similar data on ZIKV has been reported. However, maternal antibodies to dengue virus (DENV) have been described [44–47]and may facilitate enhanced disease in postnatally DENV-infected infants [46, 47].

Here, we report the results of a small study describing results from two infant macaques born to dams infected with ZIKV during the third trimester. One dam had relatively high levels of viral RNA detected in amniotic fluid near full gestation, possibly suggesting fetal infection, but no virus was detected in either infant after birth. A recent study of two infants infected with ZIKV after birth showed significant impairment of cognitive function and reduced reaction to fearful stimuli [32], which the authors interpreted as a likely consequence of infection during early infancy. Our behavioral analyses detected no indications that infant development was negatively affected by the maternal infection status. Both our study and a published study [32] performed behavioral observations on a limited number of animals and used different methods of behavioral analysis and thus cannot be directly compared. Nonetheless, behavioral data from our infants showed no direct evidence of infection nor negative consequences of infection of their dams.

Both infants in our study harbored detectable levels of anti-ZIKV neutralizing antibodies at birth that declined between one-and four-months post birth. We interpret these data to suggest these antibodies were passively acquired from the dams as opposed to mounted directly by the infants. ZIKV-binding IgG also declined after birth but remained detectable between three and five months of age, when the animals were infected. When we infected the infants with ZIKV, they exhibited low peak viremia that was rapidly cleared resulting in no evidence of infection in neurological tissues or CSF, which contrasts with published data on postnatally ZIKV-infected infant macaques [32]. ZIKV binding antibodies, likely maternal in origin, remained detectable from birth until the day of infection, possibly mediating some level of viral control. It is also possible neutralizing maternal antibodies, though undetectable in the PRNT assay at the time of infection, remained at a sufficient level to provide partial protection to the infants. In support of this possibility, infant F10, who retained detectable levels of neutralizing antibodies longer than F09, also had the lowest peak of viremia in the blood and then mounted a weaker and slower *de novo* antibody response to the virus and had less virus in lymph nodes. Alternatively, maternal antibodies may mediate protection via functions other than neutralization, such as antibody-dependent cellular cytotoxicity (ADCC) and antibody-dependent cellular phagocytosis (ADCP). Maternal antibodies with ADCC function have been detected [48]. Both infants harbored viral RNA in axillary lymph nodes at necropsy, suggesting that even a brief period of serum viremia is sufficient for tissue dissemination, which may result in consequences not tested in our study, including inflammation.

Taken together, our data suggest that infants born healthy to ZIKV infected mothers maintain a level of protection from ZIKV that dampened acute viral loads and limited tissue dissemination of the virus. We propose that passively acquired maternal antibodies might mediate a modest but important level of protection from high viremia and neurological impairment demonstrated in another NHP study of early postnatal infection.

## Author Contributions

N.J.M planned the studies and wrote the first draft. B.S., A.S., M.D., M.H.G., F.S., P.P.A., K.B., and R.V.B. conducted the experiments. N.J.M., R.P.B., K.B., K.K.A.V.R., A.A.L., M.C.B., R.V.B., S.R.P., L.L.C., A.T.P., and D.M. interpreted the studies. All authors reviewed, edited, and approved the manuscript.

## Data availability

All data is available from the corresponding author on request.

